# Sex differences in central amygdala glutamate responses to calcitonin gene-related peptide

**DOI:** 10.1101/2024.11.09.622728

**Authors:** Rebecca Lorsung, Nathan Cramer, Jason Bondoc Alipio, Yadong Ji, Sung Han, Radi Masri, Asaf Keller

## Abstract

Women are disproportionately affected by chronic pain compared to men. While societal and environmental factors contribute to this disparity, sex-based biological differences in the processing of pain are also believed to play significant roles. The central lateral nucleus of the amygdala (CeLC) is a key region for the emotional-affective dimension of pain, and a prime target for exploring sex differences in pain processing since a recent study demonstrated sex differences in CGRP actions in this region. Inputs to CeLC from the parabrachial nucleus (PB) play a causal role in aversive processing, and release both glutamate and calcitonin gene-related peptide (CGRP). CGRP is thought to play a crucial role in chronic pain by potentiating glutamatergic signaling in CeLC.

However, it is not known if this CGRP-mediated synaptic plasticity occurs similarly in males and females. Here, we tested the hypothesis that female CeLC neurons experience greater potentiation of glutamatergic signaling than males following *endogenous* CGRP exposure. Using trains of optical stimuli to evoke transient CGRP release from PB terminals in CeLC, we find that subsequent glutamatergic responses are preferentially potentiated in CeLC neurons from female mice. This potentiation was CGRP-dependent and involved a postsynaptic mechanism. This sex difference in CGRP sensitivity may explain sex differences in affective pain processing.

**Significance statement:** The central lateral nucleus of the amygdala (CeLC) receives a dense projection from parabrachial nucleus (PB) neurons that corelease calcitonin gene-related peptide (CGRP) and glutamate following aversive stimuli. This PB_CGRP_→CeLC projection plays a causal role in chronic pain. We show that endogenous CGRP release potentiates glutamate signaling in female, but not male, CeLC neurons. In the context of previous work in male CeLC, this suggests that that females are more sensitive to even transient CGRP release events. Understanding how this sex difference in CGRP sensitivity arises could enhance strategies for treating chronic pain in both women and men.

## Introduction

Women are disproportionately affected by pain, experiencing greater severity, duration, and incidence of chronic pain across many conditions (Mogil, 2009, 2012, 2021; Osborne and Davis, 2022). While societal and environmental factors can influence this sex bias (Fillingim, 2000; Bartley and Fillingim, 2013), genetic, neuroimmune and neurobiological components are thought to be involved in these sex differences (Stratton et al., 2024). Identifying mechanisms that drive sex differences in chronic pain may aid in the development of novel diagnostics and therapies to better treat women and men with these conditions.

The central lateral nucleus of the amygdala (CeLC; the “nociceptive amygdala”) is a critical center for the emotional-affective dimension of pain (Neugebauer et al., 2020). Nociceptive inputs to CeLC originate primarily from parabrachial nucleus, whose afferents form large—presumably highly efficacious— perisomatic synapses in CeLC (Delaney et al., 2007; Chou et al., 2022). CeLC integrates nociceptive and aversive inputs (Neugebauer et al., 2003; Neugebauer, 2015), and interacts with other key nodes in the pain system (Janak and Tye, 2015; Neugebauer et al., 2020). That parabrachial nucleus inputs to CeLC are causally related to persistent pain is supported by studies showing that pain-like behaviors can be suppressed by manipulating this pathway (Neugebauer, 2015; Wilson et al., 2019; Chiang et al., 2020; Raver et al., 2020; Mazzitelli et al., 2021).

Parabrachial nucleus (PB) neurons that project to CeLC express both glutamate and calcitonin gene related peptide (CGRP) (Shimada et al., 1985; Schwaber et al., 1988; Neugebauer et al., 2020). While low-frequency firing predominantly facilitates glutamate signaling, high-frequency firing of PB CGRP neurons induces the fusion of large dense core vesicles (LDCVs) (Tallent, 2008; Schöne et al., 2014; Qiu et al., 2016), releasing packaged neuropeptides including CGRP. PB neurons which express CGRP fire at these high frequencies in response to aversive input, especially in chronic pain conditions (Uddin et al., 2018; Raver et al., 2020; Smith et al., 2023). The subsequent release of CGRP upon CeLC neurons is causally related to chronic pain conditions (Han et al., 2005, 2010; Okutsu et al., 2017; Shinohara et al., 2017; Chou et al., 2022; Kang et al., 2022; Presto and Neugebauer, 2022; Allen et al., 2023; Kim et al., 2024).

Despite known sex differences in pain conditions and in pain mechanisms, including sex differences in CGRP-related pain mechanisms in humans (Labastida-Ramírez et al., 2019; de Vries Lentsch et al., 2021), essentially all data on PB and its effects on CeLC are from studies of male animals. An important exception is a demonstration that CGRP RNA levels in the CeLC are upregulated at different stages of neuropathic pain in male and female rats, and that CGRP receptor antagonist has sex-specific effects on pain behaviors (Presto and Neugebauer, 2022). Also relevant is a finding that the effects of CGRP on GABA transmission in spinal cord, and on pain behaviors, is sex-specific (Paige et al., 2022).

Here, we test the hypothesis that CGRP exerts a sex-dependent effect on glutamate signaling in the CeLC. By using a model system which combines optogenetics with patch electrophysiology, we evoke *endogenous* CGRP release from PB_CGRP_ terminals in the CeLC *in vitro*. By relying on endogenous release of CGRP, rather than exogenous application of CGRP, we minimize the risk of off target effects by more closely mimicking physiologic release of neuropeptides. We first validate that single optic stimulation of channel rhodopsin (ChR2) expressing PB_CGRP_ terminals induces glutamate release in the CeLC, while high frequency stimulation is required to induce neuropeptide release *in vitro*. We then test the effect of endogenously released CGRP on glutamate signaling. We predicted that CeLC glutamate signaling is potentiated by CGRP signaling, in line with previous studies in male rodents (Han et al., 2010; Okutsu et al., 2017), in both sexes, but with a greater magnitude of potentiation in female neurons.

## Methods

### Animals

All procedures adhered to the Guide for the Care and Use of Laboratory Animals and approved by the Institutional Animal Care and Use Committee at the University of Maryland School of Medicine. We used 25 CGRP (calcitonin-gene-related peptide)-CRE heterozygous mice (13 female, 12 male) that were bred in house from male B6.Cg-Calca^tm1.1(cre/EGFP)Rpa^/J (stock #033168) x female C57BL/6J mice (strain #000664). Breeding pairs were obtained from The Jackson Laboratory. Offspring were weaned at postnatal day (PD)21 and housed two to five per cage in single-sex groups. Food and water were available ad libitum, and lights were maintained on a 12/12 h light/dark cycle. Two males (M1-2) and two females (F1-2) were used for fiber photometry experiments. The remaining mice were used for *in vitro* electrophysiology, where 1-3 neurons were recorded in each mouse from 1-3 CeLC slices.

### Virus injection

We anesthetized the animals with isoflurane and placed them in a stereotaxic frame. Either left or right PBN (−5.2 mm AP, ±1.5 mm ML, −2.9 mm DV) was targeted via a small craniotomy (∼1-2 mm). Only the right PBN was targeted in LDCV photometry recordings. We injected 0.5 μL of adeno-associated virus generated by the University of Maryland School of Medicine’s Viral Vector Core – Baltimore, Maryland; AAV5-DIO-ChR2-eYFP, OR 0.25 μL AAV_DJ_-DIO-CYbSEP2 co-injected with AAV5-DIO-ChR2-eYFP. CybSEP2 is a presynaptic pH-sensitive presynaptic sensor which is trafficked by LDCVs, and which undergoes a shift in fluorescence upon LDCV fusion and neuropeptide release (Kim et al., 2024). Viruses were injected using a MICRO2T SMARTouch™ controller and Nanoliter202 injector head (World Precision Instruments) at a flow rate of 100 nL/min. The pipette was left in place for 10 min before being slowly retracted over 5–10 min. Mice were given Rimadyl for postoperative analgesia. Injection sites were verified by visually confirming robust eYFP fluorescence in the external PBN.

### In vitro slice electrophysiology

We anesthetized adult mice (2 – 12 months old) generated live brain slices from adult mice and generated 300µm thick coronal sections through the central nucleus of the amygdala using a modified slice collection method as described in (Ting et al., 2014) and our prior studies. For recordings, we placed slices in a submersion chamber continuously perfused (2 mL/min) with artificial cerebrospinal fluid (ACSF): 119 mM NaCl, 2.5 mM KCl, 1.2 mM NaH_2_PO_4_, 2.4 mM NaHCO_3_, 12.5 mM glucose, 2 mM MgSO_4_·7H_2_O, and 2 mM CaCl_2_·2H_2_O. ACSF was adjusted to a pH of 7.4, mOsm of 305, and bubbled with carbogen (95% O_2_ and 5% CO_2_) throughout use. We obtained whole-cell voltage-clamp recordings (−70 mV) from the capsular region of the CeLC using borosilicate pipettes with an impedence of 4-6 MΩ and containing: 130 mM cesium methanesulfonate, 10 mM HEPES, 1 mM magnesium chloride, 2.5 mM ATP-Mg, 0.5 mM EGTA, 0.2 mM GTP-Tris, 5 mM QX-314, and 2% biocytin (pH of 7.3, 285 mOsm). Excitatory postsynaptic currents (EPSCs) were optically evoked by whole field illumination at 470 nm (Lambda LS light source, Sutter Instrument) and maximum power of 1.4 mW. Optical stimulation parameters, both high frequency stimulation (10 or 20 Hz, 3 ms pulse duration) and single/paired exposures (3 ms pulse duration, 100 ms interval), were controlled by a SmartShutter system (Sutter Instrument). We monitored series resistance by measuring the current evoked by a −5 mV square pulse at ∼ 20s intervals. Evoked oEPSC amplitudes were quantified using Clampfit 11.2 (Molecular Devices).

### In vitro LDCV quantification

Acute brain sections containing the CeLC were collected as above (see “In vitro slice electrophysiology”) from adult mice previously injected with AAV_DJ_-DIO-CYbSEP2 (Kim et al., 2024) and AAV5-DIO-ChR2-eYFP in the ipsilateral parabrachial nucleus. A fiber optic probe (400 μM diameter, 0.39 NA; RWD Life Sciences) was placed over the visually identified fluorescent afferents from PB within CeLC to record LDCV sensor transients evoked by high frequency optical stimulation of PB_CGRP_ fibers in the CeLC at 470 nm (CoolLED). Sensor transients were recorded through the fiber optic probe connected to a RZX10 LUX fiber photometry processor running Synapse software (Tucker-Davis Technologies) through a Doric mini cube (Doric Lenses). Fiber photometry LED power was calibrated to 15 µW using a digital optical power meter (Thor Labs).

We analyzed the data using customized Python scripts adapted from Tucker-Davis Technologies templates which calculated relative changes in fluorescence. Changes in sensor fluorescence were calculated by subtracting the scaled isosbestic signal (405 nm) from the sensor fluorescence (465 nm). Event related changes in sensor fluorescence were converted to ΔF/F using the 5 second window prior to each stimulation as baseline. The area under the curve (AUC) for the average response was calculated for each mouse using the AUC analysis function in GraphPad Prism.

### Experimental Design and Statistical Analysis

Statistical tests were conducted using Prism 10 (GraphPad), and sample size was determined using G*Power software suite (Heinrich-Heine, Universität Düsseldorf). Parametric tests were used when appropriate assumptions were met; otherwise, we used nonparametric tests. Specific statistical tests are detailed in Table 1. Unless describing a time course, baseline vs post optic comparisons are shown as the average in the 90 seconds before and 90 seconds after optic tetanus delivery. Averages described in the text are formatted as mean ± SD unless otherwise stated. All figures were designed using a combination of Prism 10 (GraphPad) and Inkscape 1.3.2. Atlas images for Figure 5A were adapted from (Franklin and Paxinos, 2008), and accessed via a web based tool (https://labs.gaidi.ca/mouse-brain-atlas/).

**Table 1:** Statistics for showing the corresponding figure number, animals, metric, comparisons being made, test statistic, medians or means, sample size, and p values.

| Figure | Animals | Metric | Comparison | Test Statistic | Mean | Sample size | p Value |
| --- | --- | --- | --- | --- | --- | --- | --- |
| 2C | F3-13, M3-7 | Normalized oEPSC amplitude | CGRP vs Baseline within sex | Female paired t test ($t=4.971$, $df=15$)<br>Male paired t-test ($t=0.5583$, $df=7$) | Female: Baseline 1.00, CGRP 1.36<br>Male: Baseline 1.00, CGRP 1.06 | 16 female neurons, 8 male neurons | Female: $p=0.0002$, Male: $p=0.59$ |
| 2D | F3-13, M3-7 | Normalized oEPSC amplitude | Sex, Time, Sex x Time | RM Mixed-effects analysis<br>Sex ($F(1, 314) = 9.794$),<br>Time ($F(6.229, 139.7) = 2.629$),<br>Sxt ($F(14, 314) = 2.080$)<br>Dunnet's multiple comparison test performed vs baseline average ($\alpha = 0.05$) | Timepoint from left to right, 0.05 Hz sampling (m, f):<br>Baseline<br>1) 0.86, 0.97<br>2) 0.99, 1.18<br>3) 1.00, 1.05<br>4) 0.98, 0.99<br>5) 0.90, 0.94<br>6) 0.84, 1.12<br>7) 0.856, 1.01<br>8) 1.03, 1.00<br>9) 1.07, 0.924<br>10) 1.34, 0.91<br>CGRP<br>1) 1.26, 1.53<br>2) 0.89, 1.45<br>3) 0.97, 1.32<br>4) 0.89, 1.17<br>5) 0.74, 1.00 | 8 male neurons, 15 female neurons | Sex: $p < 0.01$<br>Time: $p = 0.02$<br>SxT: $p = 0.01$ |
| 2E | F3-13, M3-12 | Normalized baseline holding current | After vs before within sex | Female paired t-test ($t=2.267$, $df=14$),<br>Male Wilcoxon test ($W=54.00$) | Female: Before 1.00, After - 1.52<br>Male: Before 1.00, After 3.22 | 15 female neurons, 15 male neurons | Female: $p=0.04$, Male: $p = 0.14$ |
| 2F | M8-12 | Normalized oEPSC amplitude | CGRP vs baseline | Paired t test ($t=0.5971$, $df=6$) | Male: Baseline 1.00, CGRP 0.90 | 7 male neurons | |
| 2G | M8-12 | Normalized oEPSC amplitude | Time | RM Mixed effects analysis<br>Time ($F(2.203, 12.75) = 0.6122$), Dunnet's multiple comparison test performed vs baseline average ($\alpha = 0.05$) | Timepoint from left to right, 0.033 Hz sampling:<br>Baseline<br>1) 1.00<br>2) 0.98<br>3) 1.00<br>4) 0.86<br>5) 1.38<br>6) 1.14<br>7) 1.03<br>8) 0.87<br>9) 0.91<br>10) 1.08<br>CGRP<br>1) 0.92<br>2) 0.90<br>3) 0.91<br>4) 0.95<br>5) 0.73 | 7 male neurons | P=0.57 |
| 2H | F3-13, M3-12 | Response magnitude (>20% for increased/decreased) | Male vs female CeLC neurons | Fisher's exact test | N/A | 15 male neurons, 16 female neurons | P=0.0032 |
| 3A | F3-4, 6-8, 9-10 | oEPSC amplitude normalized to baseline | Between treatments | Paired t test ($t=0.5296, df=7$) | CGRP<sub>1</sub>: 1.533<br>CGRP<sub>2</sub>: 1.472 | 8 female neurons | P=0.61 |
| 3B | F4, 6, 8-9, 11-12 | oEPSC amplitude normalized to baseline | Between treatments | Paired t test ($t=2.960, df=7$) | CGRP<sub>1</sub>: 1.545<br>CGRP<sub>2</sub> + Antagonist: 1.149 | 8 female neurons | P=0.02 |
| 3C | F3-4, 6, 8-9, 12-13 | Normalized paired pulse ratio (PPR) | Between treatments | Paired t test ($t=1.558, df=9$) | Baseline: 1<br>CGRP: 0.847 | 10 female neurons | P=0.15 |
| 4A | F3-13 | Normalized potentiation amplitude | Right vs left CeLC (female only) | Unpaired t test ($t=0.7812, df=13$) | Left: 1.492, Right: 1.360 | 7 left, 8 right CeLC neurons | P=0.45 |
| 4B | F3-13 | oEPSC baseline amplitude | Right vs left CeLC (female only) | Unpaired t test (t=0.008863, df=17) | Left: -24.22, Right: -24.15 | 9 left, 10 right CeLC neurons | p>0.99 |
| 4C | F3-13 | Baseline Paired pulse ratio (PPR) | Right vs left CeLC (female only) | Unpaired t test (t=0.3136, df=11) | Left: 0.59, Right: 0.55 | 6 left, 7 right CeLC neurons | P=0.76 |
| 4D | F3-9,11-13 | Mouse age | Responding vs nonresponding female neurons | Unpaired t test (t=0.2681, df=13) | Non responder: 32.17, responder: 30.54 | 6 non responder and 9 responder mice | P=0.79 |
| 4E | F3-9,11-13 | Virus incubation | Responding vs nonresponding female neurons | Mann-Whitney test (U = 15) | Non responder median: 7.14, responder median: 12.57 | 6 non responder and 9 responder mice | P=0.17 |
| 4F | F3-9,11-13 | Access resistance | Responding vs nonresponding female neurons | Mann-Whitney test (U=20) | Non responder median: 0.83, responder median: 0.42 | 5 non responder and 11 neurons | P=0.44 |
| 4G | F3-13 | Baseline holding current | Responding vs nonresponding female neurons | Mann-Whitney test (U=29) | Non responder median: -37.00 responder median: -38.00 | 6 non responder and 10 neurons | P=0.96 |
| 4H | F3-9,11-13 | Membrane resistance | Responding vs nonresponding female neurons | Unpaired t test (t=0.8766, df=13) | Non responder: 947.8, responder median: 685.1 | 6 non responder and 9 responder mice | P=0.4 |
| 4I | F3-9,11-13 | Cell capacitance | Responding vs nonresponding female neurons | Unpaired t test (t=0.02697, df=13) | Non responder: 107.1, responder median: 106.1 | 6 non responder and 9 responder mice | P=0.98 |
| 5B | F3-13, M3-12 | Proportion patched CeLC neurons with evoked<br>oEPSCs | Male vs female | Fisher's exact test | N/A | 21 male and 23 female CeLC neurons<br>patched | P=0.449 |
| 5C | F3-13,<br>M3-12 | oEPSC<br>amplitude | Male vs<br>female | Unpaired t test (<br>t=0.05990, df=32) | Male: -24.57,<br>female: -24.19 | 15 male<br>and 19<br>female<br>neurons | P=0.95 |
| 5D | F3-13,<br>M3-12 | Baseline<br>holding<br>current | Male vs<br>female | Mann-Whitney<br>test (U=96) | Male median: -<br>50.0, Female<br>median: -37.90 | 14 male<br>and 18<br>female<br>neurons | P=0.27 |
| 5E | F3-13,<br>M3-12 | Access<br>resistance | Male vs<br>female | Mann-Whitney<br>test (U=104) | Male median:<br>0.56, Female<br>median: 0.79 | 14 male<br>and 16<br>female<br>neurons | P=0.76 |
| 5F | F3-9,11-13, M3-12 | Membrane<br>resistance | Male vs<br>female | Unpaired t test<br>(t=0.4094, df=30) | Male: 760.1,<br>female: 847.9 | 15 male<br>and 17<br>female<br>neurons | P=0.69 |
| 5G | F3-9,11-13, M3-12 | Cell<br>capacitance | Male vs<br>female | Unpaired t test<br>(t=0.4610, df=30) | Male: 113.1,<br>female: 847.9 | 15 male<br>and 17<br>female<br>neurons | P=0.65 |

## Results

### High frequency optical stimulation of PB_CGRP_ terminals induces LDCV release in the CeLC

Parabrachial nucleus (PB) neurons that express calcitonin gene-related peptide (CGRP; PB_CGRP_) co-release glutamate and CGRP. To evoke glutamate and CGRP release from these terminals in CeLC, we injected a Cre-dependent channel rhodopsin (ChR2) virus into the PB of CGRP-Cre mice (Fig. 1A). The CeLC receives dense input from PB CGRP neurons (Fig. 1B), whose cell bodies densely localize in the lateral parabrachial nucleus (LPB, Fig. 1C). We first recorded optically-induced excitatory postsynaptic currents (oEPSCs) from ipsilateral CeLC neurons evoked by a single pulse of 470 nm light (3 msec duration, 0.05 Hz) (Fig.1). Application of AMPA receptor antagonist (CNQX, 20 μM) suppressed oEPSCs, indicating these oEPSCs are dependent on glutamate signaling. Similar suppression was observed in 4 of 4 neurons. While these single optic stimuli reliably induced oEPSCs in CeLC neurons, high frequency stimulus trains (10 Hz, 5 sec), that mimic firing patterns of PB neurons during noxious stimulation in normal and chronic pain states (Uddin et al., 2018; Raver et al., 2020; Smith et al., 2023), resulted in rapid suppression in oEPSC amplitudes recorded in CeLC neurons from both male and female mice (Fig. 1E, 11 female neurons; 8 male neurons). This suggests that the strength of glutamatergic excitation of CeLC neurons postsynaptic to PB afferents rapidly diminishes during sustained activity.

**Figure 1.**
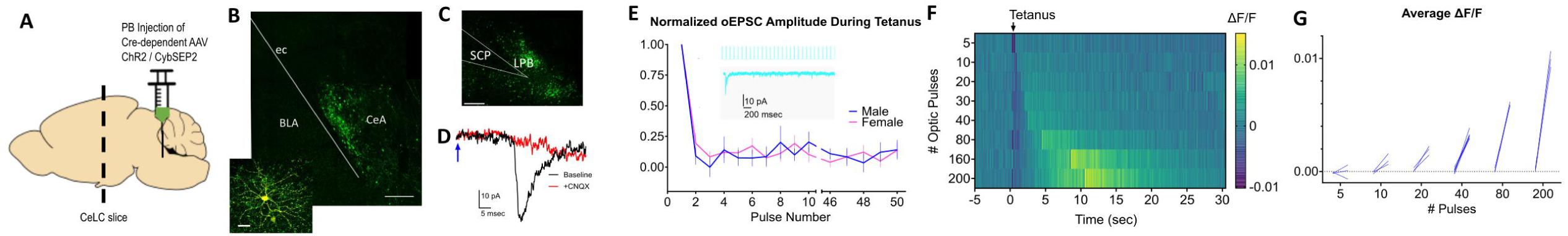
Low frequency optic stimulation of PB_CGRP_ terminals in CeLC induces glutamate release, while high frequency optic stimulation induces neuropeptide release. (A) AAV5-DIO-ChR2-eYFP OR AAV5-DIO-ChR2-eYFP AND AAVDJ-DIO-CYbSEP2 virus was injected into the left or right lateral parabrachial of CGRP-Cre mice. CeLC slices were collected for either fiber photometry or voltage clamp patch electrophysiology. (B) Photomicrographs showing PB_CGRP_ EYFP+ terminals in ipsilateral central amygdala (CeLC). Scale bar = 200 μm. A biocytin filled CeLC neuron shown in inset. Scale bar = 20 μm. (C) An injection site, with dense eYFP+ PB_CGRP_ neuronal cell bodies located in lateral parabrachial (LPB). Scale bar = 200 μm. SCP = superior cerebellar peduncle. (D) A single optic stimulus induces an optically evoked EPSC (oEPSC) at baseline (black trace) which is suppressed by 20 μM CNQX (red). Blue arrow indicates delivery of single optic stimulus (3 msec duration). (E) Rapid reduction in oEPSC amplitudes during delivery of a high frequency optic tetanus. Example stimulus train and response in CeLC neuron shown in inset (cyan). (F) Heat map depicting the change in CybSEP2 fluorescence (ΔF/F) in response to different durations of 20 Hz stimulus train of ChR2-expressing PB_CGRP_ inputs in a male CeLC slice. Three trials averaged per row. (G) Quantification of average fluorescence in 5 seconds pre and post stimulus for trials shown in F.

While high frequency activity causes a drop in glutamatergic drive, we reasoned that these same stimulus trains would be sufficient to drive release of CGRP from large dense core vesicles (LDCVs) (Tallent, 2008; Schöne et al., 2014; Qiu et al., 2016). To test this, we directly measured neuropeptide release from PB_CGRP_ afferents which co-express channelrhodopsin2 (ChR2) and the LDCV sensor CybSEP2. CybSEP2 is a pH sensitive sensor transported by LDCVs, which undergoes a shift in fluorescence upon LDCV fusion and neuropeptide release (Kim et al., 2024). We co-injected Cre-dependent CybSEP2 and Cre-dependent ChR2 into the PB of CGRP-cre mice (Fig. 1A) and performed fiber photometry in slices from ipsilateral CeLC to monitor LDCV release. Figure 1F depicts a heat map obtained from recordings where the duration of the stimulus train increased from 250 ms (5 pulses at 20 Hz) to 10 seconds (200 pulses at 20 Hz). As evidenced by the corresponding ΔF/F (Fig. 1F) and quantified as area under the curve (AUC, Fig. 1G), short duration stimulus trains failed to produce detectable changes in sensor fluorescence. However, increasing the stimulus duration progressively increased recorded fluorescent intensity (Fig. 1G). Similar responses were seen in slices from male and female mice (n = 2). In contrast, mice injected with a single virus, either CybSEP2 (n=1) or ChR2 (n=1), demonstrated no such increase. These data indicate that high frequency optic stimulation is sufficient to induce PB_CGRP_ LDCV fusion, and subsequent neuropeptide release, *in vitro* in the CeLC.

### PB glutamatergic signal is potentiated in females following PB endogenous CGRP release

We took advantage of this ability to evoke PB neuropeptide release to determine if endogenous CGRP release potentiates PB_CGRP_->CeLC glutamatergic synapses. To measure the amplitude of this glutamatergic signaling we recorded responses to single pulses of blue light (3 msec duration, 0.05 Hz) to induce ChR2-mediated glutamatergic oEPSCs in postsynaptic CeLC neurons (Fig. 2). After establishing the baseline oEPSC amplitude, we used high frequency optical stimulation (10 Hz, 5 sec) to evoke neuropeptide release in the CeLC and then remeasured the glutamatergic response to single pulse stimulation. Figure 2A depicts oEPSC recorded from a CeLC neuron from a female mouse, before and after CGRP release, demonstrating a 44% increase in the amplitude of the glutamatergic response after the tetanus stimulation. In contrast, Figure 2B shows that the same procedure failed to potentiate oEPSCs in a CeLC neuron from a male. Group data for female and male mice are quantified in Figure 2C, where CeLC neurons from female mice displayed an average potentiation of 136%±29% (p=2×10^−3^, N=16 cells from 11 female mice) following high frequency tetanus, while male CeLC neurons showed no net potentiation, with an average post-optic tetanus magnitude of 106%±29% (p=0.59, N=8 cells from 5 male mice). The time course of this response is shown in Figure 2D where oEPSCs were significantly potentiated only in female mice (RM two-way mixed-effects sex F (1, 314) = 9.794, p<0.01; time F (6, 229) = 2.629, p=0.02; sex x time F (14, 314) = 2.080, p=0.01), and up to 60 s after the tetanus (Dunnet’s multiple comparison test, p<0.05). Additionally, in female CeLC neurons, there was a small reduction of 1.5±2.6 pA in holding current (p=0.04; 15 neurons, 10 mice) in the 5 seconds following high frequency optic tetanus (Fig. 2E). This suggests there was a temporary change in membrane ion conductance induced by the high frequency optic tetanus. In males there was no net change in holding current (p=0.16; 15 neurons, 9 mice, Fig. 2E).

**Figure 2.**
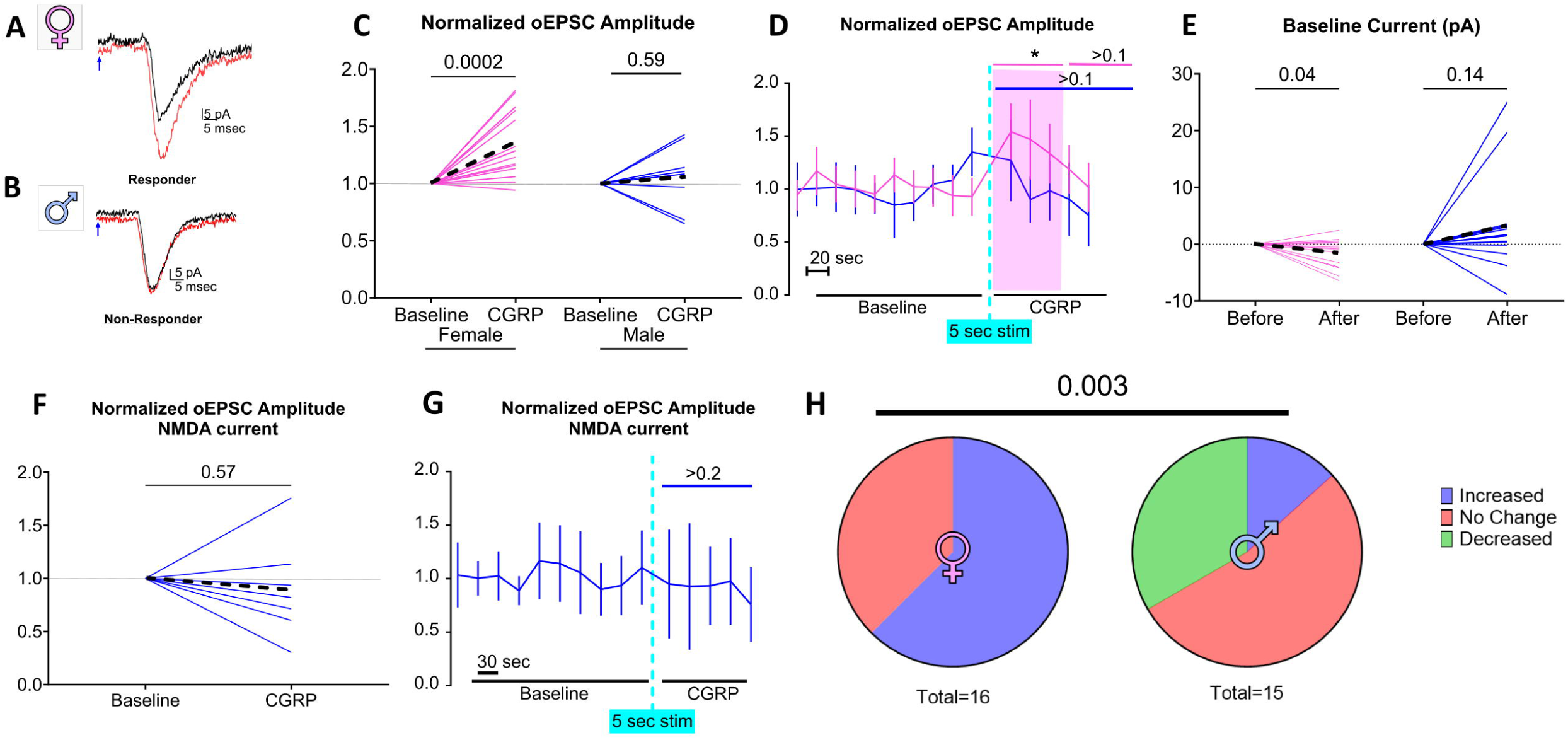
Optic stimulation of PB CGRP induces a transient, CGRP-dependent potentiation of glutamate sensitivity, preferentially in females. PB_CGRP_ neurotransmitter release was induced via single (glutamate) or high frequency (CGRP) optic tetanus. oEPSCs evoked by single-pulse optic stimulation from CeLC neurons from a female(**A**) and a male (**B**) neuron recorded at baseline (black traces) and post PB CGRP release (red traces). Note the potentiation in the female, but not the male neuron. Blue arrows indicate delivery of single optic stimulus (3 msec duration). (**C**) Amplitudes of optically evoked PB_CGRP_ glutamate responses (oEPSC) in the 80 second period following high frequency optic in female and male CeLC neurons. Dashed lines depict averages. (**D**) Amplitude of oEPSC, normalized to baseline, transiently potentiated for ∼60 seconds in female CeLC neurons (pink). (**E**) A shift in baseline holding current specifically in females accompanied this high frequency optic stimulus. Using a higher frequency optical train to enhance PB CGRP release (20 Hz, 10 sec) and recording in 0 mg solution to enhance NMDA currents failed to evoke oEPSC potentiation (**F,G**). (**H**) Significant difference in the effects of endogenous CGRP release on female and male neurons. Increases and decreases were defined as >20% change in oEPSC amplitudes.

To test whether the observed sex differences in potentiation arise from a higher threshold for potentiation in CeLC neurons from males versus females, we use a higher frequency, longer optical tetanus to encourage greater endogenous CGRP release (20 Hz, 10 sec) while recording from a population of CeLC neurons patched in separate male mice (N= 7 neurons, 5 mice). In addition, as previous studies demonstrated that CGRP acts through a NMDA-dependent mechanism in the CeLC (Han et al., 2010; Okutsu et al., 2017), we amplified the signal from recruited NMDA channels by excluding Mg^2+^ in the ACSF. Even under these conditions there was no net potentiation in male CeLC neurons, as post-optic tetanus oEPSC amplitude was 89%±46% of baseline (p=0.57, Fig. 2F). Figure 2G depicts the time course of this response (RM one-way mixed-effect of time F(2.203, 12.75)=0.6122, p=0.57).

Across both experiments, only 13% of recorded male CeLC neurons (2/15 neurons, 10 mice) were potentiated (>20% elevation of oEPSC amplitude) following high frequency stimulation. In contrast, potentiation occurred in 62% of female CeLC neurons (10/16 neurons, 11 mice). Figure 2H depicts the responses across all recorded neurons, where male and female CeLC neurons respond differently to high frequency optic stimulation (p=0.003).

Potentiation in females was reproducible within a neuron; once response amplitudes returned to baseline, a similar level of potentiation (p=0.61 compared to the previous potentiation) was achieved by a second delivery of high frequency optic stimulation (N=8 neurons, 5 mice; Fig. 3A). Re-potentiation in female CeLC neurons was suppressed by pre-application of 1 μM of the CGRP receptor antagonist (CGRP_8-37_, p=0.02), suggesting that CGRP signaling contributes to this increase in PB_CGRP_->CeLC glutamate signaling (N= 8 neurons, 6 mice, Fig. 3B).

**Figure 3.**
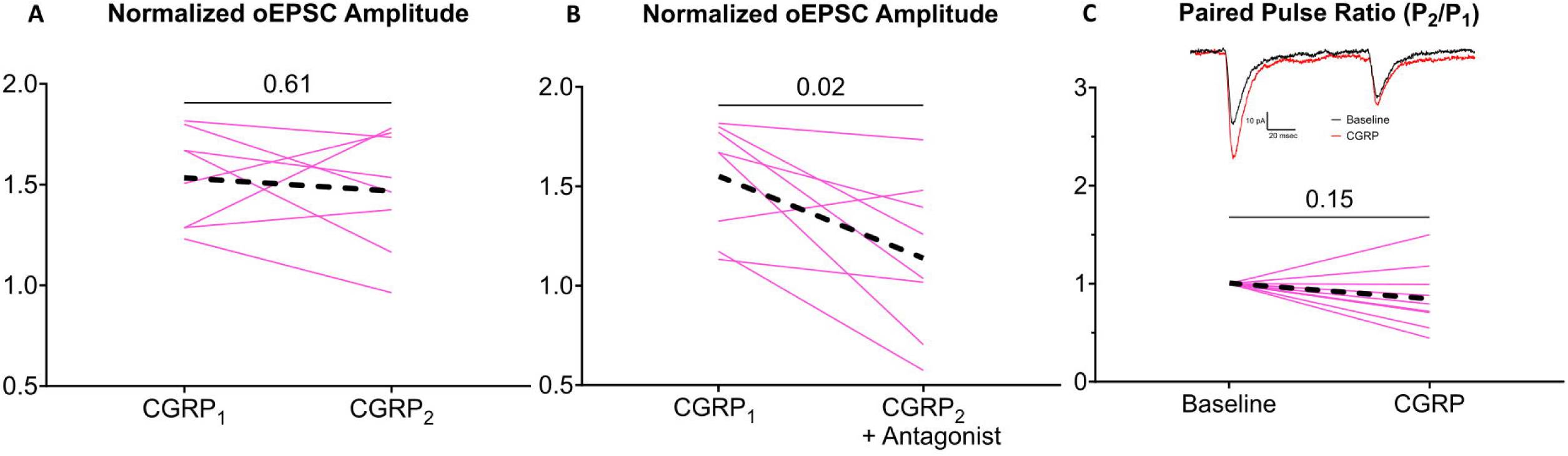
Potentiation is CGRP-dependent and influences presynaptic release probability in females. (**A**) Potentiation of PB_CGRP_ glutamate signaling in female neurons is reproducible, as a second PB CGRP release event (CGRP_2_) induces a similar level of potentiation as an initial PB CGRP release event (CGRP_1_). (**B**) This potentiation is CGRP-dependent as evidenced by its suppression by 1 μM CGRP8-37, CGRP receptor antagonist. (**C**) The potentiation is not associated in with a change in paired pulse ratios, indicating that it does not involve presynaptic mechanisms. Black dashed lines indicate group averages. Inset shows traces of paired pulses (100 msec interstimulus interval) at baseline (black) and after high frequency optic tetanus (red).

### Effect of high Frequency optic tetanus on presynaptic release probability and passive membrane properties

We tested whether CGRP dependent potentiation in female neurons is driven by a presynaptic mechanism by comparing oEPSC responses to paired pulse stimulation and calculated a paired pulse ratio (PPR), the ratio of the amplitude of the second pulse divided by the amplitude of the first pulse; changes in paired pulse ratio implicate the involvement of a presynaptic mechanism (Manabe et al., 1993; Debanne et al., 1996; Dobrunz and Stevens, 1997; Kim and Alger, 2001). A PPR above one is associated with a low probability of vesicle release (i.e., “weaker” synapses), whereas a PPR below one is associated with a high probability of release. Example paired-pulse oEPSCs are shown in Figure 3C (inset). We compared PPR before and after optic tetanus delivery. At baseline, the oEPSC PPR was less than 1 in all female CeLC neurons, suggesting these are high release probability synapses. Female CeLC neurons (10 neurons, 7 mice) did not exhibit a change in PPR following CGRP-mediated potentiation (p=0.15, Fig. 3C). This suggests that CGRP acts postsynaptically.

To test if CGRP mediated potentiation reflects postsynaptic mechanisms, we compared holding current before and after potentiation. While there was a small reduction in holding current in female CeLC neurons immediately following the high frequency optic tetanus (Fig. 2E), there was no change in holding current PB CGRP release compared to baseline in either male (p=0.16; 14 neurons, 9 mice) or female (p=0.21; 16 neurons, 11 mice) CeLC neurons. This suggests that while female CeLC neurons are transiently depolarized in response to high frequency optic tetanus, the potentiated glutamatergic response outlasts the change in driving potential evoked by the tetanus itself. There was no change in series resistance following PB CGRP release in either male (p=0.6; 14 neurons, 9 mice) or female (p=0.62; 16 neurons, 11 mice) CeLC neurons. Therefore, the effects of endogenous CGRP release upon CeLC neurons likely reflect a localized change to the post synapse, and not a more far-reaching alteration to the overall intrinsic membrane properties of the CeLC neurons.

### CGRP-dependent potentiation and input strength are consistent between hemispheres and across the A-P axis of the CeLC

Lateralization in the function of CeLC has been reported in several preclinical models of pain (Carrasquillo and Gereau, 2008; Ji and Neugebauer, 2009; Allen et al., 2023). Therefore, we examined whether lateralization of PB_CGRP_ optic tetanus-evoked potentiation occurs in female neurons. There was no difference (p=0.45; 15 neurons, 10 mice) in potentiation magnitude between left and right CeLC (Fig. 4A). Additionally, there was no difference in baseline oEPSC amplitude (p=0.99; 18 neurons, 12 mice) nor in baseline paired pulse ratio (p=0.76; 18 neurons, 12 mice) between recordings from the left and right CeLC (Fig. 4B-C). These findings indicate that there is no lateralization in PB_CGRP_->CeLC glutamatergic input strength or synaptic release probability in females.

**Figure 4.**
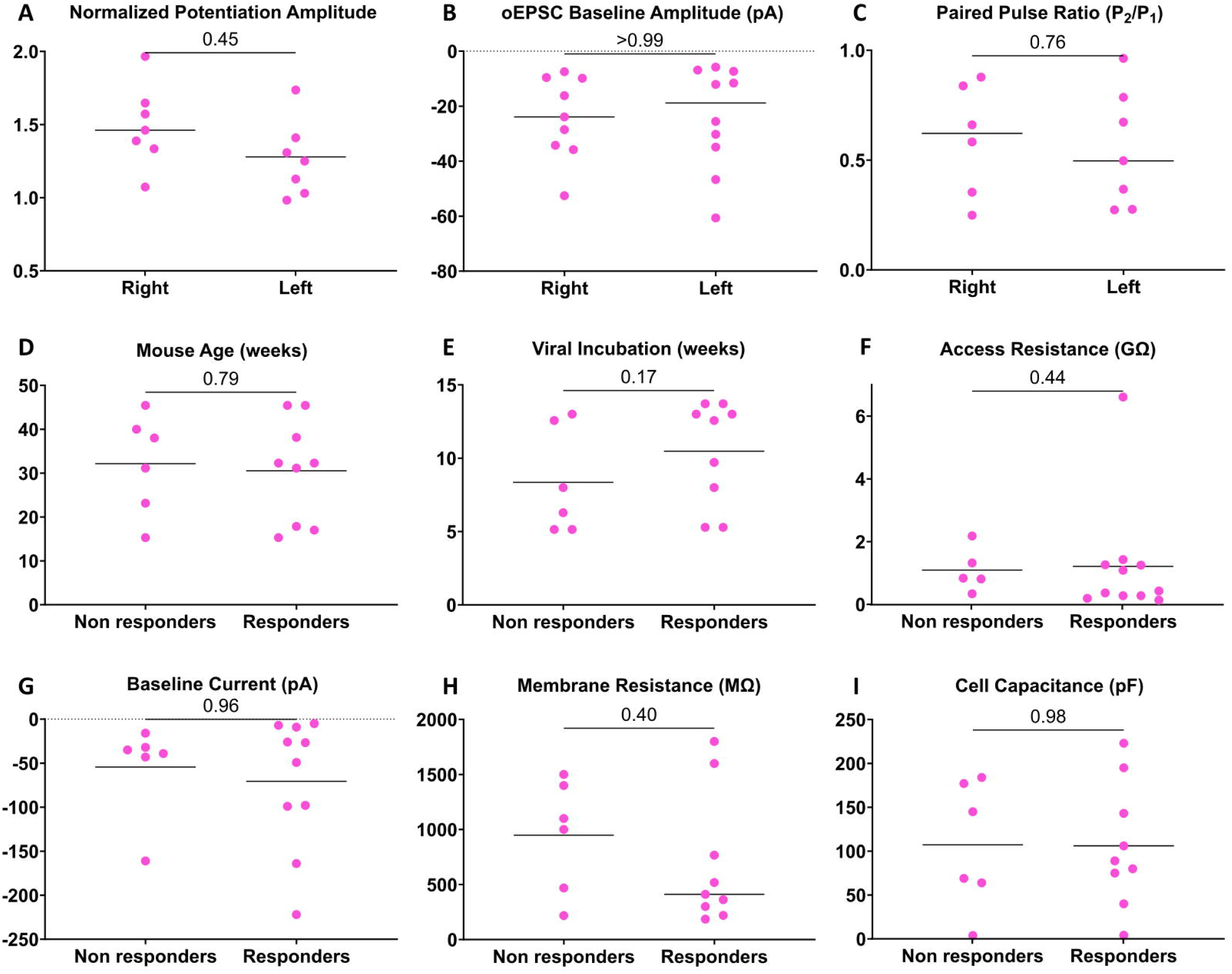
CGRP-dependent potentiation and input strength is not lateralized, and response to CGRP is not due to experimental variables. All data from female CeLC neurons. Potentiation is indistinguishable in neurons in the left or right CeLC (**A**). Two measures of PB_CGRP_ glutamatergic input strength, oEPSC baseline amplitudes (**B**) and paired pulse ratios at baseline (**C**) also show no lateralization. Potential experimental sources of heterogeneity are not correlated with potentiation – animal age (**D**); virus incubation time (**E**) do not differ between responder neurons (>20% potentiation) and non-responder neurons. Similarly, access resistance (**F**), baseline current (**G**), membrane resistance (H) or cell capacitance (**I**) show no difference between responders and non-responders.

CeLC may differ functionally not only between hemispheres, but also across the anterior-posterior axis. For example, optogenetic stimulation of PB_CGRP_ inputs to the anterior and posterior CeLC induces different behaviors; stimulating PB_CGRP_ in posterior CeLC inputs induces freezing, while stimulating PB_CGRP_ in anterior CeLC predominantly induces changes in respiration and vasoconstriction (Bowen et al., 2020). Therefore, we compared the sensitivity of the female CeLC across the A-P axis to glutamatergic potentiation following PB CGRP release. In female CeLC neurons where oEPSCs were detected in response to optic stimulation of PB_CGRP_ terminals (direct PB_CGRP_ glutamatergic input), widefield images of the CeLC were acquired following patching. The neuronal anterior-posterior position relative to bregma was then determined using a stereotaxic atlas (Franklin and Paxinos, 2008). The location of recorded neurons was not related to PB CGRP-driven glutamate potentiation (r=0.26, p=0.36; 14 neurons, 8 mice), baseline PB glutamate amplitude (r=0.16, p=0.53; 17 neurons, 10 mice), or baseline PB glutamate paired pulse ratio (r=0.10, p=0.74; 13 neurons, 8 mice) in females. This suggests both PB CGRP and PB glutamate exert comparable effects on glutamatergic input from PB across anterior and posterior CeLC in females.

To determine if failure of neurons to potentiate results from experimental variables, we compared several experimental properties between female CeLC neurons that exhibited CGRP-dependent potentiation (>20%) and those that did not. We compared animal age (p=0.79; 16 neurons, 11 mice, Fig. 4D), viral incubation times (p=0.17; 16 neurons, 11 mice, Fig. 4E), and access resistance (p=0.44; 16 neurons, 11 mice, Fig. 4F). Experimental parameters did not differ between potentiated and non-potentiated neurons. We also compared passive membrane properties: baseline holding current (p=0.96; 16 neurons, 11 mice, Fig. 4G), membrane resistance (p=0.40; 15 neurons, 10 mice, Fig. 4H) and cell capacitance (p=0.98; 15 neurons, 10 mice, Fig. 4I); variations in cell capacitance and resistance in particular can reflect differences in the electrical “control” of a patch and resulting resolution. Passive membrane properties did not differ between potentiated and non-potentiated neurons.

### Parabrachial glutamate inputs to central amygdala show no sex differences

We considered the possibility that the sex differences in synaptic potentiation are related to sex differences in the underlying glutamatergic component of the PB_CGRP_->CeLC signaling pathway. Figure 5A displays the location of CeLC neurons receiving glutamatergic PB_CGRP_ input. There were no sex differences in the proportion of CeLC neurons receiving PB_CGRP_ input (p=0.44; 23 female neurons, 11 mice; 21 male neurons, 10 mice, Fig. 5B); over 75% of both male and female CeLC neurons exhibited evoked oEPSCs following optic stimulation of PB_CGRP_ glutamate release. There was also no sex difference in baseline oEPSC amplitudes (p=0.95, 15 male neurons, 10 mice; 19 female neurons, 11 mice, Fig. 5C). This suggests that the strength of PB_CGRP_->CeLC synapses is similar in males and females.

**Figure 5.**
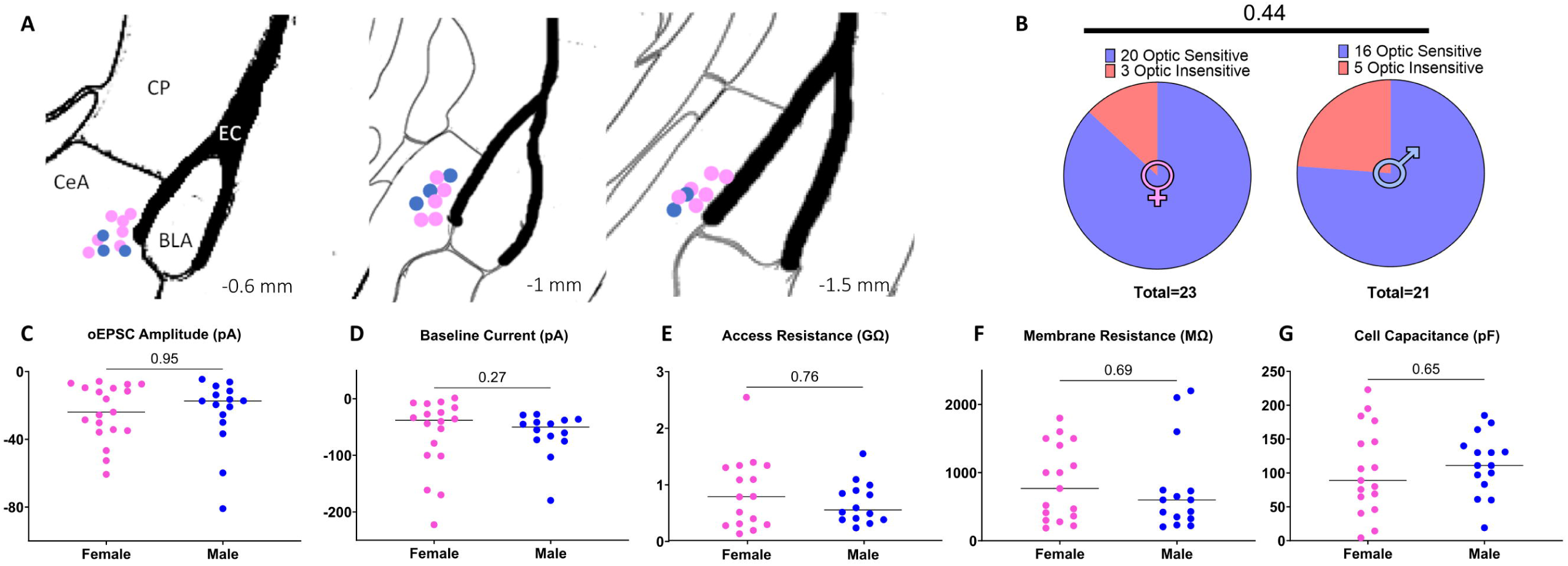
Male and female CeLC neurons recording locations and membrane properties. (**A**) Location of male (blue) and female (pink) CeLC neurons that received direct PB_CGRP_ glutamatergic input recorded in the lateral capsule of the CeLC across a number of coronal slices. Positions indicated relative to bregma. (**B**) There was no sex difference in the proportion of recorded CeLC neurons that received direct PB_CGRP_ glutamatergic input. (**C**) There was no sex difference in oEPSC amplitudes at baseline, a metric of input strength. Male and female CeLC neurons also had similar baseline currents **(D**), access resistance (**E**), membrane resistance (**F**), or membrane capacitance (**G**).

The CeLC contains a variety of GABAergic neuronal classes (Schiess et al., 1999; Xu et al., 2003; Lopez De Armentia and Sah, 2004; Chieng et al., 2006; Amano et al., 2012; Lu et al., 2015; Wilson et al., 2019). The sex differences in response to endogenous CGRP release may relate to sex difference in the subpopulations of CeLC neurons receiving direct PB_CGRP_ input. While functionally distinct central amygdala subclasses are primarily defined by their firing pattern (Schiess et al., 1999; Lopez De Armentia and Sah, 2004; Chieng et al., 2006; Amano et al., 2012; Wilson et al., 2019), subtle differences in intrinsic neuronal properties between these classes have also been observed, particularly resting membrane potential (Schiess et al., 1999; Chieng et al., 2006). We observed no sex differences in intrinsic neuron properties including holding current (p=0.27; 14 male neurons, 10 mice; 18 female neurons, 11 mice, Fig. 5D), which is the current required to maintain a constant membrane potential. Similarly there was no sex difference in access resistance (p=0.76, 13 male neurons, 9 mice; 17 female neurons, 12 mice, Fig. 5E) membrane resistance (p=0.69; 15 male neurons, 10 mice; 17 female neurons, 12 mice, Fig. 5F), nor cell capacitance (p=0.65; 15 male neurons, 10 mice; 17 female neurons, 12 mice Fig. 5G).

## Discussion

### Endogenous CGRP release

The central amygdala (CeLC) densely expresses a number of neuropeptides and neuropeptide receptors (Neugebauer et al., 2020). Among these is CGRP, which can increase CeLC neuronal activity in response to aversive processing, especially in chronic pain conditions (Han et al., 2005, 2010; Shinohara et al., 2017; Neugebauer et al., 2020). Previous studies used exogenously applied CGRP to reveal these effects of CGRP on CeLC neuronal functions. However, these results are confounded by the fact that both the physiological concentration and clearance rate of CGRP in CeLC are unknown. As a result, it is not known if exogenously applied CGRP mimics physiological conditions. Here, we circumvented these limitations by endogenously inducing neuropeptide release in the CeLC using optogenetics to drive parabrachial CGRP-expressing (PB_CGRP_) activity.

High frequency firing is necessary for neuropeptide release (Schöne et al., 2014; Qiu et al., 2016). We reasoned that high frequency firing of PB_CGRP_ neurons that occurs in response to aversive stimuli (Uddin et al., 2018; Smith et al., 2023) would provide the presynaptic activity required for CGRP release. Indeed, simulating this firing by driving PB_CGRP_ neurons optogenetically resulted in release of large dense core vesicles (LDCVs) from PB_CGRP_ neurons in the central amygdala, as evidenced by imaging of LDCV sensor signals (Fig. 1F). Further, these stimuli trains resulted in potentiation of CeLC PB_CGRP_ glutamate response (Fig. 2C). That this potentiation was suppressed by a CGRP antagonist confirms it involves activation of CGRP receptors (Fig. 3B). These findings indicate that *endogenous* release of CGRP can potentiate the activity of CeLC neurons.

### Sex differences

There was a sex difference in CeLC neuron response to endogenous CGRP release. Most (62%) CeLC neurons in females exhibit transient, CGRP-dependent potentiation to PBCGRP glutamatergic signaling after high frequency stimulation of PBCGRP inputs, compared to 13% of male neurons. In contrast, previous studies (Han et al., 2010; Okutsu et al., 2017) describe a similar level of glutamatergic potentiation (∼140%) in *male* CeLC neurons from both rats and mice with exogenous CGRP. This discrepancy might be due to lower CGRP sensitivity in males, or insufficient CGRP release to activate receptors in male CeLC neurons. This hypothesis is supported by the kinetics of CGRP-driven potentiation in male CeLC, which required >7 minutes of continuous CGRP exposure (Han et al., 2010; Okutsu et al., 2017), compared to our findings in female CeLC, where 5 seconds of CGRP release were sufficient to induce glutamate potentiation. No potentiation was observed in males even with increased PB_CGRP_ stimulation. A sex difference in CGRP sensitivity is consistent with behavioral studies demonstrating enhanced CGRP-dependent pain and anxiety-like behaviors in females (Avona et al., 2019; Paige et al., 2022), and clinical data showing greater efficacy of CGRP-targeting therapies in women (Porreca et al., 2024). CGRP antagonism in CeLC also blocks affective pain behavior exclusively in females (Presto and Neugebauer, 2022).

Sex differences in the effects of CGRP may arise from several mechanisms. CGRP clearance may be more active in males. CGRP degradation remain a field of active study (Russo and Hay, 2023), with a variety of enzymes implicated in CGRP clearance, including but not limited to matrix metalloproteinase 2 (Fernandez-Patron et al., 2000) and neutral endopeptidase (Katayama et al., 1991; Davies et al., 1992; McDowell et al., 1997). Sex differences in clearance mechanisms have been observed in some (Zhao et al., 2011; Howe et al., 2019; Omori et al., 2020), but not all studies (Reuveni et al., 2017; Bronisz et al., 2023).

Additionally, males may release less CGRP in the CeLC. PB neurons are equally activated by noxious stimuli in males and females (Smith et al., 2023), and no sex differences in LDCV release was observed *in vitro* using a novel fluorescent indicator (observation from Kim et al., 2024). However, it is possible that males either release fewer LDCVs in response to equivalent PB neuron activity, or CGRP is less densely expressed among packaged neuropeptides within LDCVs. This has not been tested, although we did not observe potentiation in males even upon increasing the hypothetical “dose” of CeLC CGRP by escalating PB_CGRP_ stimulation. Finally, considering the relatively long timescale for potentiation initiation described in males (Han et al., 2010; Okutsu et al., 2017), it is possible that bath application of CGRP exerts its potentiating effects via signaling from an intermediate cell. CGRP receptors are robustly expressed on a variety of glial cells, including endothelial cells (Crossman et al., 1990), where they act to promote vasodilation (Brain et al., 1985). CGRP’s effects on other glial types remains to be determined.

There were no sex differences in PB_CGRP_ glutamatergic input. The majority of CeLC neurons in both sexes received PB_CGRP_ glutamatergic input, in line with previous literature describing robust PB input to the “nociceptive amygdala” (Han et al., 2010; Okutsu et al., 2017). Additionally, input strength was similar between sexes, with no sex difference in the amplitude of optically evoked PB_CGRP_ glutamate currents. This suggests that low-frequency activity of PB activity, dominated by glutamate signaling, has similar effects on CeLC neurons of both sexes. However, high-frequency PB firing during chronic pain (Helassa et al., 2018; Uddin et al., 2018; Raver et al., 2020) likely shifts signaling to neuropeptide predominance. Thus, functional consequences of a sex difference in CGRP’s effect in the CeLC may be restricted to aversive conditions.

### Synaptic mechanisms

Short term potentiation describes transient (seconds to minutes) potentiation of glutamatergic synapses (Fioravante and Regehr, 2011), as described here in female CeLC neurons following endogenous CGRP release. A postsynaptic mechanism has been implicated in male CeLC, where CGRP engages PKA-dependent NMDA receptor recruitment to the PBCGRP synapse (Han et al., 2010; Okutsu et al., 2017). Our findings are consistent with these data, as we observed no change in paired pulse ratio in female CeLC neurons following CGRP release (Fig. 3C).

### Potentiation kinetics

Endogenous CGRP release transiently potentiates glutamatergic input for (∼60 seconds following a 5 second stimulus train), whereas exogenous CGRP maintains potentiation at least throughout CGRP application (Han et al., 2010; Okutsu et al., 2017), and up to 30 minutes afterward (Okutsu et al., 2017). This difference in the potentiation kinetics may be due to clearance dynamics. Endogenous CGRP release results in physiological CGRP concentrations, which are subject to clearance and diffusion away from CGRP receptors. It is possible that exogenous application of CGRP overwhelms endogenous CGRP clearance, allowing for continuous activation of CGRP receptors, and thus maintained potentiation of glutamate signaling.

### Response heterogeneity

Following CGRP release, 62% of female CeLC neurons showed potentiated glutamate signaling. This heterogeneity was not explained by experimental factors such as animal age, viral incubation time, passive membrane properties, or patch access. Different CeLC regions are implicated in various functions, such as anterior/posterior CeLC activation inducing divergent behavioral/physiological responses (Bowen et al., 2020), and pain lateralizing to the right CeLC (Neugebauer and Li, 2003; Han and Neugebauer, 2004; Carrasquillo and Gereau, 2008; Ji and Neugebauer, 2009; Sadler et al., 2017; Allen et al., 2023). However, CeLC neuron location could not explain this observed heterogeneity; we found no differences in the magnitude of CGRP-dependent potentiation, nor PB_CGRP_ glutamatergic input strength, between hemispheres or across the anterior-posterior axis.

Heterogeneity in female CeLC responses may reflect the molecular diversity of CeLC GABAergic neurons. While both PKCδ and somatostatin (SST)-expressing CeLC neurons - subpopulations of CeLC GABAergic neurons - receive PB glutamatergic inputs (Wilson et al., 2019) and express CGRP receptors (Han et al., 2015; Chou et al., 2022), these populations play opposing roles in nociception (Wilson et al., 2019). It is, therefore, possible that these populations have different sensitivities to endogenous CGRP signaling. This could arise from differences in the subcellular locations of CGRP receptors relative to the site of CGRP release. Differences in the location of PB_CGRP_ terminal contacts in PKCδ and SST cells support this possibility (Shimada et al., 1989; Ye and Veinante, 2019), where perisomatic CGRP^+^ terminals closely surround PKCδ-expressing CeLC neurons, but rarely SST-expressing neurons. However, these studies did not specifically investigate the location of CGRP receptors. CGRP-induced changes in proximal synapses, such as those observed on PKCδ-expressing CeLC neurons, would be more easily resolved due to superior voltage clamp control at synapses closer to the neuronal cell body (Spruston et al., 1993). Alternatively, there could be differences in signaling downstream from the CGRP receptor; while PKA-dependent (Gα_s_) processes are confirmed in male neurons (Han et al., 2010; Okutsu et al., 2017), both Gα_i_ and Gα_q_ processes have been implicated in male cardiac tissue, cultured astrocytes, and immortalized cell lines (Walker et al., 2010).

Even transient aversive PB hyperactivity induces PB neuropeptide release in the CeLC (Kim et al., 2024). Our findings suggest that the female nociceptive amygdala is particularly vulnerable to excitation during these CGRP release events, whereas males require sustained elevated CGRP to induce similar potentiation (Han et al., 2010; Okutsu et al., 2017). This sex difference in CGRP sensitivity may underlie sex differences in affective pain processing, with females showing heightened vulnerability to pain and its related affective sequalae (Mogil, 2009, 2012, 2021; Osborne and Davis, 2022).

## Acknowledgments

This work was supported by National Institutes of Health–National Institute of Neurological Disorders and Stroke Grants R01NS099245, R01NS069568, R01NS127827 and Fellowship F31NS134126 (to RL).

Dr. Jason Alipio performed work on this project while in the Department of Neurobiology. He is currently affiliated with the Center for Regenerative Medicine, Massachusetts General Hospital, Boston, MA, USA. 02118

